# ampir: an R package for fast genome-wide prediction of antimicrobial peptides

**DOI:** 10.1101/2020.05.07.082412

**Authors:** Legana C.H.W Fingerhut, David J. Miller, Jan M. Strugnell, Norelle L. Daly, Ira R. Cooke

## Abstract

**Summary:** Antimicrobial peptides (AMPs) are key components of the innate immune system that protect against pathogens, regulate the microbiome, and are promising targets for pharmaceutical research. Computational tools based on machine learning have the potential to aid discovery of genes encoding novel AMPs but existing approaches are not designed for genome-wide scans. To facilitate such genome-wide discovery of AMPs we developed a fast and accurate AMP classification framework, ampir. ampir is designed for high throughput, integrates well with existing bioinformatics pipelines, and has much higher classification accuracy than existing methods when applied to whole genome data.

**Availability and Implementation:** ampir is implemented primarily in R with core feature calculation methods written in C++. Release versions are available via CRAN and work on all major operating systems. The development version is maintained at https://github.com/legana/ampir

**Supplementary information:** Supplementary data are available at https://github.com/legana/amp_pub

## Introduction

Antimicrobial peptides (AMPs) are produced by most forms of life to combat microbial pathogens including bacteria, viruses, fungi, and protists. Their potent activity has led to strong interest in these molecules as targets for pharmaceutical research. Additionally, AMPs are key regulators of the microbiome, and understanding this role is emerging as an important focus of AMP research (Thaiss *et al.*, 2016).

Despite intense interest in AMPs, the genes that encode them remain difficult to detect. They evolve rapidly, driven by positive selection coupled with high rates of gene gain and loss (Hanson *et al.*, 2019), and this, combined with the small size of mature AMP peptides (10-50 amino acids), makes them difficult to detect through homology based approaches alone. A promising alternative approach to AMP detection is the use of supervised machine learning based on physicochemical properties. Many AMP predictors have been developed using this approach, with recent examples including iAMPpred (Meher *et al.*, 2017), AmPEP (Bhadra *et al.*, 2018), AMP Scanner (Veltri *et al.*, 2018), and dbAMP (Jhong *et al.*, 2019). In principle, software of this type could be used to enhance AMP discovery by helping to identify candidates from whole transcriptome or genome data (see for example Yoo et al., 2014).

Despite several AMP predictors being available, we found that they have several shortcomings when used to perform a whole genome scan. Firstly, the user interfaces of most existing predictors are not designed to facilitate high throughput or integrate with other tools into automated pipelines. With few exceptions (e.g. AmPEP), they are available primarily through web applications that lack a programming interface and have restrictions on the number of sequences that can be analysed in one batch. A second, and perhaps more critical issue affecting existing AMP prediction approaches is that they are trained and evaluated using datasets that do not resemble typical whole genome data, leading to poor prediction accuracy in genome scans (Figure 1). There are two main aspects to this dataset mismatching (detailed summaries are available in sections S1 and S2). One issue is that most existing predictors are trained and evaluated on mature peptides whereas in a genome-scanning context it is much more likely that researchers will be working with full-length precursor protein sequences. The second issue is that whole genome data typically contains a very small proportion (usually less than 1%) of AMPs but evaluation datasets used by existing approaches are either balanced (50% AMPs) or nearly balanced leading to unrealistically low estimates of the false positive rate.

**Figure 1:**
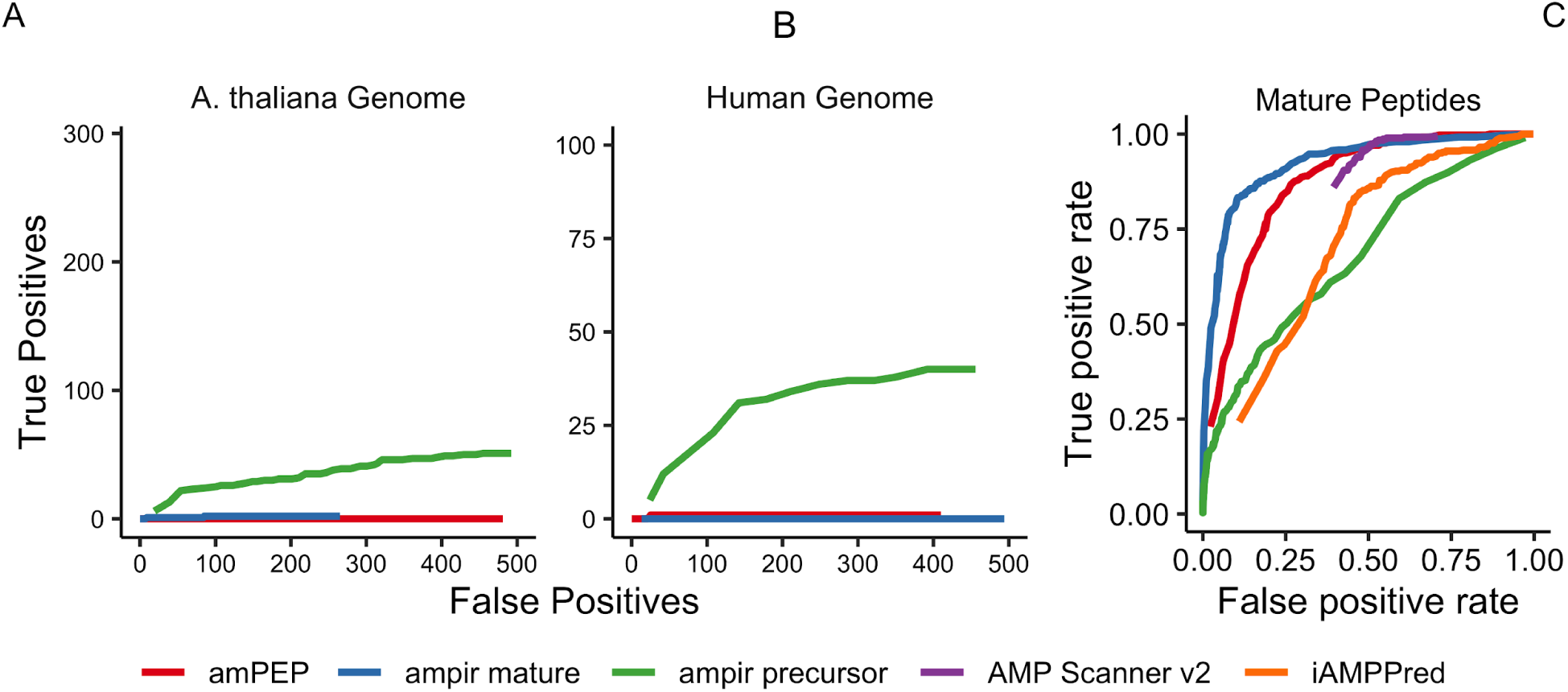
Performance of ampir compared with three existing AMP prediction models: iAMPpred, AmPEP and AMP Scanner v2. Results for iAMPpred are not shown for parts A and B because it was impractical to run on large numbers of sequences. Parts A and B are scaled so that the limits of the y-axis show the full complement of known AMPs in each genome (291 for *A. thaliana*, 101 for Human), and the limits of the x-axis are restricted to emphasise behaviour in the low false positive regime because this is most relevant in whole genome scans. Part C is a receiver operating characteristic (ROC) curve based on a 20% fraction of the ampir_mature dataset that was reserved for evaluation.

## Implementation and benchmarking

ampir includes two built-in models, one trained on mature peptide data (ampir_mature) which is comparable with previous approaches and another trained on full-length precursor data (ampir_precursor). Complete details of the design of these classifiers is provided online at https://github.com/legana/amp_pub and in the supplementary information (section S3). Both models were built as support vector machines with a radial kernel. The full-length precursor model was trained using unbalanced data (9% AMPs) to provide an accurate representation of feature space while using class weights to avoid over emphasising the majority class. Model hyperparameters (sigma, C) were optimised based on 10-fold cross validation on 80% of the data with the remaining 20% reserved for performance testing.

Most existing predictors performed very well (AUROC 0.86-1.0) when evaluated with a widely used benchmark dataset from Xiao et al. (2013), containing mature peptides and short proteins, as this had a similar composition to training data used (section S2). Arguably a more realistic task is to distinguish mature peptides with AMP activity from other mature peptides that do not share this role. The ampir_mature model was trained with this in mind and performed better than other predictors when evaluated against a benchmark dataset composed entirely of mature peptides (Figure 1C, Table S1).

We also performed a genome-scanning benchmark using the well-annotated genomes of Human and *Arabidopsis thaliana* (Figures 1A, 1B). While this is a more realistic benchmark for whole genome scanning performance, it is also much more challenging because AMPs comprise less than 1% of the protein sequences in these datasets (0.14% in Human, 0.74% in *A. thaliana*) so even a highly accurate predictor will produce many false positives. Here, the ampir_precursor model was the only classifier capable of predicting an appreciable proportion of true AMPs, while restricting the number of false positives to less than 500 (Figure 1A & B). This demonstrates that ampir is capable of generating a result set that is greatly enriched in AMPs from a whole genome scan, and would therefore be valuable as the first stage in a screening pipeline for AMP discovery.

### ampir as a framework

In addition to providing built-in classifiers for mature peptide sequences or full-length precursor proteins, ampir provides a framework to allow researchers to easily build custom models and use them for fast genome-wide prediction. Specifically, ampir provides computationally efficient methods (implemented in C++ with multicore support) for calculating features commonly used in AMP prediction including physicochemical properties (Meher *et al.*, 2017; Bhadra *et al.*, 2018) and Chou’s pseudo amino acid composition (Xiao *et al.*, 2013) and uses the Peptides package (Osorio et al., 2015) to calculate all other physicochemical properties commonly used in AMP prediction (Meher et al., 2017; Bhadra et al., 2018). The underlying model training and prediction capabilities are provided by the caret package (Kuhn, 2008). A complete scan of all 77,000 proteins in the Human proteome with ampir can be completed in under four minutes on a desktop PC (four cores).

We anticipate that in genome-scanning contexts such custom models will be especially important since they allow for optimisation (a) within a restricted taxonomic range, or (b) with restricted or biased input data (e.g. only secreted proteins). Researchers can take full advantage of the caret framework to optimise for these contexts on the basis of training data, feature selection and underlying machine learning approach. The resulting models can then be provided directly to ampir for prediction.

