## Supplementary Information for "ampir: an R package for fast genome-wide prediction of antimicrobial peptides"

Legana C.H.W Fingerhut[1,2,3*], David J. Miller[1,2,3], Jan M. Strugnell[4], Norelle L. Daly[1,5] and Ira R. Cooke[1,2]

1 Centre for Tropical Bioinformatics and Molecular Biology, College of Public Health, Medical and Veterinary Sciences, James Cook University, Townsville, Qld, 4811, Australia.

2 Department of Molecular and Cell Biology, James Cook University, Townsville, Queensland, Australia

3 ARC Centre of Excellence for Coral Reef Studies, Australian National University, Canberra, ACT, Australia

4 Centre for Sustainable Tropical Fisheries and Aquaculture, College of Science and Engineering, James Cook University, Townsville, Qld, 4811, Australia.

5 Centre for Molecular Therapeutics, Australian Institute of Tropical Health and Medicine, James Cook University, Cairns, Qld, 4870, Australia.

This document provides a summary of methods used to collect and curate training data, train models and perform benchmarks. In the interests of reproducibility and transparency all data and code is provided, along with detailed descriptions at https://github.com/legana/amp_pub

### Table of Contents

Figure S1: Sequence length distributions in publicly available AMP databases. ... 3
A total of 182 SwissProt proteins and 52 dbAMP proteins with sequence lengths greater than 300 amino acids were excluded due to constraints on the x-axis... 3
Figure S2: Sequence length distribution of SwissProt entries classified according to whether they were mature peptides, precursor proteins or unknown... 4
*Section S2: Summary of existing prediction software* ... 4
Figure S3: Length distribution of positive (AMP) and negative (non-AMP) training sequences for AmPEP, Amp scanner v2 and the mature peptide benchmark data from Xiao et al. ... 5
*Section S3: ampir default classifiers* ... 5
Precursor Model Training Data ... 5
Mature Peptide Model Training Data ... 6
Feature Calculation and Selection ... 7
Model Training and Tuning ... 7
*Section S4 Model performance evaluation on various benchmark sets* ... 7
Table S1: Performance of ampir and other AMP predictors on three different test sets ... 7
Table S2: Performance of ampir and other AMP predictors on ampir's training data ... 8
Figure S4: Performance of various AMP predictors in classifying whole proteome data for Human and Arabidopsis thaliana. Performance is shown as a balance of sensitivity and precision (top row) and as a ROC curve (second row). ... 10
*Section S5 Running time* ... 10
Table S2: Run time performance on the Human Proteome ... 11
Figure S5: Performance of ampir as a function of core count when running predict_amps() on a dataset of 77,000 proteins ... 11
*References:* ... 12

### Section S1: Summary of antimicrobial peptide databases

We benchmarked the performance of ampir against three existing AMP prediction models: iAMPpred (Meher et al. 2017), AmPEP (Bhadra et al. 2018) and AMP Scanner (Veltri et al. 2018). The training data used for these models and ampir was derived from one or more of the following AMP databases:

> APD: The Antimicrobial Peptide Database was accessed in March 2020 via the query interface (http://aps.unmc.edu/AP/database/query_input.php) and contained a total of 3,177 AMPs.
>
> DRAMP: This database lists both natural and synthetic AMPs. We downloaded natural AMPs in March 2020 via http://dramp.cpu-bioinfor.org/browse/NaturalData.php which contained 4394 sequences.
>
> dbAMP: The latest release of dbAMP is from 06/2019 and was downloaded from http://140.138.77.240/~dbamp/download.php. It contains 4213 experimentally verified natural AMPs (synthetic AMPs were removed).
>
> UniProt: AMPs were downloaded from UniProt (https://www.uniprot.org/) on 14-April-2020 using the search term: “keyword:Antimicrobial [KW-0929]”. This included 3221 reviewed and 19288 unreviewed proteins. The reviewed fraction of this dataset is called “SwissProt”.

One of the most striking differences between AMP databases becomes clear simply by looking at the length distributions (Figure S1). The APD, DRAMP and dbAMP databases emphasise short peptides (mostly < 50 amino acids (AA) whereas SwissProt includes many more longer proteins. The pronounced double peak in the distribution of SwissProt proteins is likely due to the fact that it contains both full length precursors (peak around 80 AA) and mature peptides (peak around 25 AA). This is investigated further below.

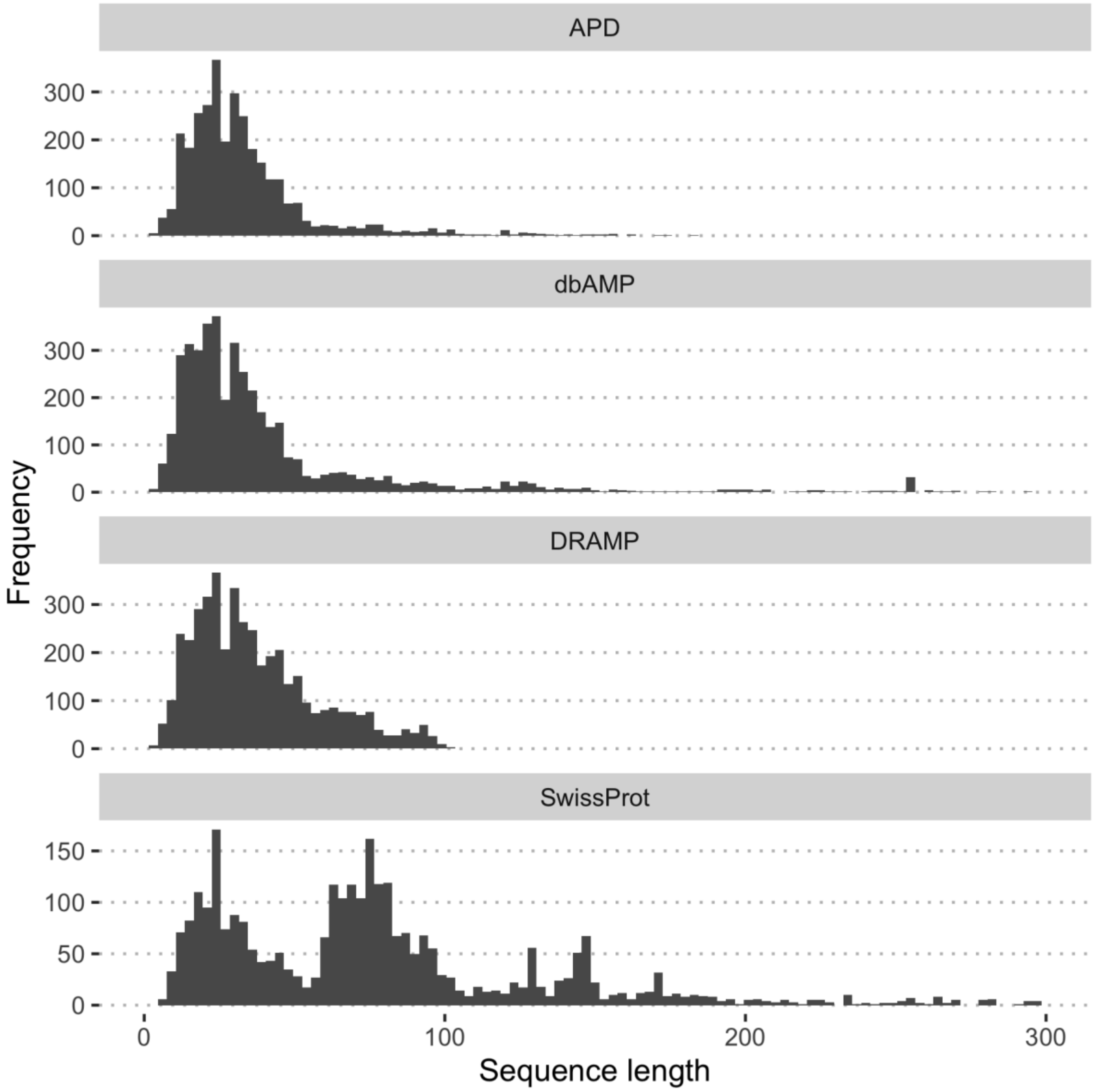


**Figure S1: Sequence length distributions in publicly available AMP databases.**

**A total of 182 SwissProt proteins and 52 dbAMP proteins with sequence lengths greater than 300 amino acids were excluded due to constraints on the x-axis.**

The SwissProt database includes detailed information on post-translational processing for many proteins, potentially allowing for clear differentiation between mature peptides and their precursors. More specifically, some protein entries in SwissProt include a Peptide field, which we used to calculate the length and position of mature peptides, in the protein. In cases where the entire SwissProt entry is a mature peptide this field is present and shows that the peptide length is the same as the total length. We used this to identify mature peptides. In cases where the peptide field was present, but did not correspond to the full length of the protein we classified that entry as a full length precursor. In some cases the Peptide field was absent entirely and we classified those as unknown. Plotting the length distributions of proteins in these categories (Figure S2) confirms that the double-peak shown in Figure S1 most likely represents the distinction between mature peptides and full length precursors. We also found that almost all (706/806) proteins identified as precursors based on the peptide field had well defined signal peptide sequences indicating that they fit the classical model of an AMP precursor protein (Zasloff, 2002).

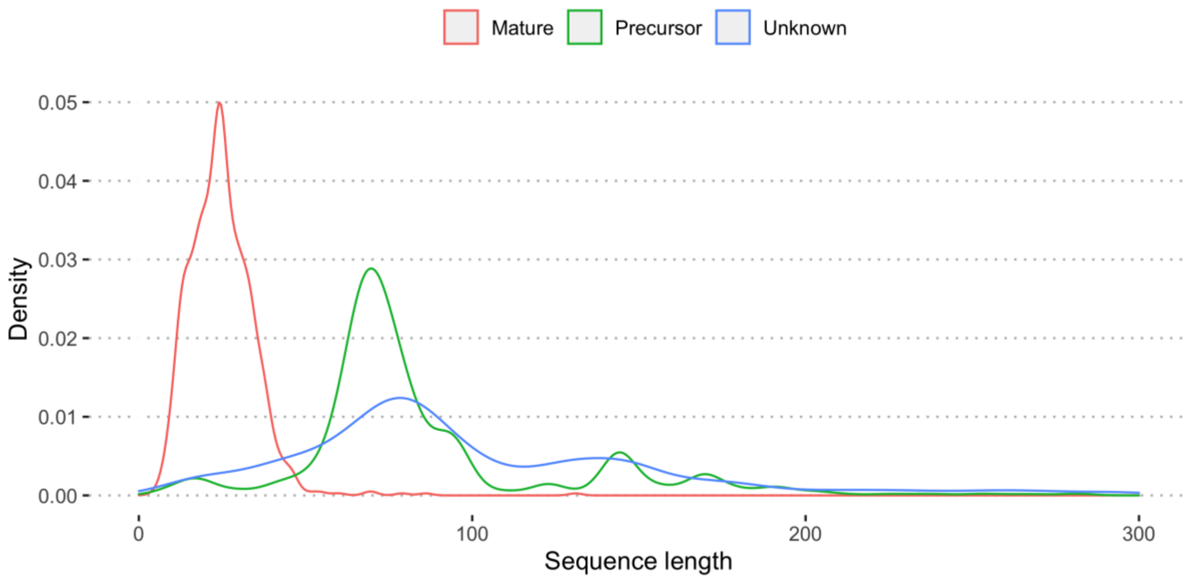


**Figure S2: Sequence length distribution of SwissProt entries classified according to whether they were mature peptides, precursor proteins or unknown.**

### Section S2: Summary of existing prediction software

iAMPpred: This predictor attempts to classify peptides specifically into antibacterial, antiviral and antifungal. It does this by including three separate SVM based predictors, one for each AMP type. Since our purpose was simply to classify a peptide as AMP or non-AMP we took the maximum value of the three available predictions as the probability that the peptide was an AMP of some type. Details of the datasets used for training these three predictors are provided in (Meher et al., 2017) and included the AMP database APD which we show above in Figure S1 is largely composed of mature peptides. Sequence lengths were restricted to between 10 and 100 which means that some full length precursors are likely included. On this basis we expect that iAMPpred includes a mix of mature peptides and full length precursors. For the purpose of benchmarking all iAMPpred results were obtained from the iAMPpred webserver http://cabgrid.res.in:8080/amppred/ in April 2020.

AmPEP: The complete training and test datasets for AmPep are available for download. These clearly show (Figure S3) that the positive dataset is largely composed of peptides in the correct length range to be mature peptides and that this differs strongly from the length distribution of the background data.

AMP Scanner v2: The complete training, test and benchmarking datasets used to build the classifier were obtained from the AMP Scanner website https://www.dveltri.com/ascan/v2/about.html. The length distribution of training sequences is shown in Figure S3 and shows a clear emphasis on mature peptides. In this case the non-AMP length distribution reflects the fact that the authors matched it (through random substring sampling) to that of the positive (AMP) dataset.

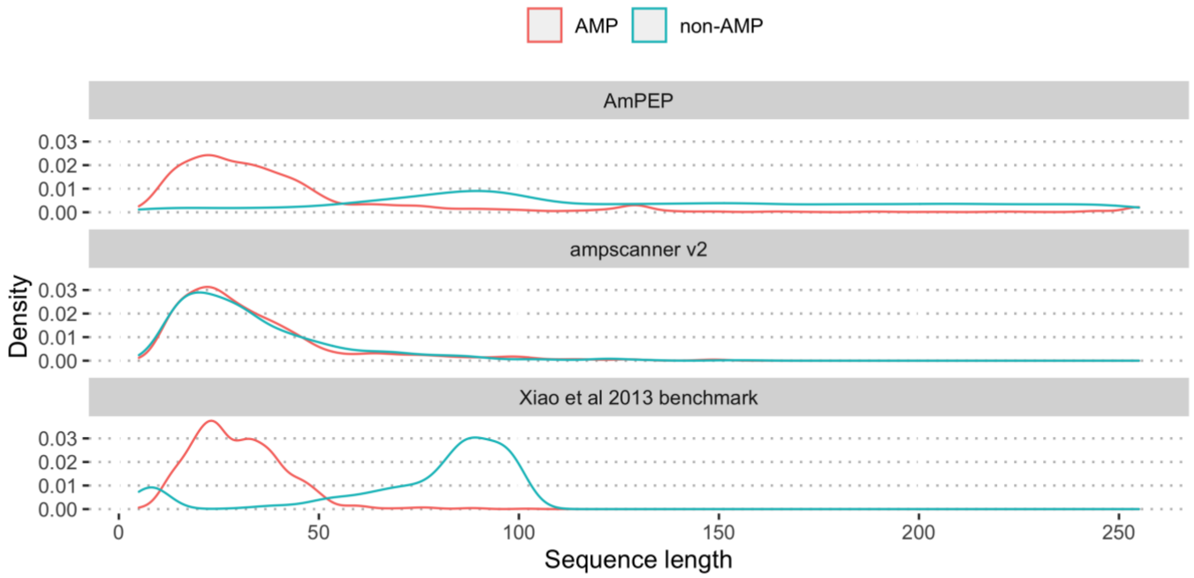


**Figure S3: Length distribution of positive (AMP) and negative (non-AMP) training sequences for AmPEP, Amp scanner v2 and the mature peptide benchmark data from Xiao et al.**

### Section S3: ampir default classifiers

Although ampir allows users to supply their own custom models it also comes with two models designed to satisfy major use cases, namely to classify mature peptides and precursor proteins respectively. The model building process for each of these is described below:

#### <u>Precursor Model Training Data</u>

For the precursor protein model we selected positive (AMP) cases using the UniProt database (including both reviewed and unreviewed proteins) as a starting point. This was then refined as follows to identify positive (AMP-precursor) cases:

1. Remove any mature peptide entries based on the Peptide field
2. Exclude unreviewed entries unless they appeared in APD, DRAMP or dbAMP
3. Remove short sequences (<50 AA) since these might represent mature peptides imported from APD, DRAMP or dbAMP
4. Remove very large proteins (>500 AA) since these are likely to have very different physicochemical properties and are not amenable to prediction by this method
5. Remove identical sequences and sequences with non-standard amino acids
6. Cluster sequences to 90% identity using CD-HIT (Fu *et al.*, 2012)

This resulted in a total of 1,483 distinct positive cases.

Since precursor proteins encompass a wide range of potential sequences in SwissProt the number of available sequences for the background dataset is much larger (over 300k) than the number of known AMP precursor proteins. This imbalance is also broadly reflective of a typical genome-scan where AMPs would be expected to comprise around 1% or less of the genome. We found it was possible to take advantage of this abundance of background data by using class weights to avoid over emphasising the negative class. Although we did not perform an exhaustive search over different class imbalances we found that an AMP:non-AMP ratio of 1:10 achieved similar results to larger ratios while reducing fitting times and model memory usage. In all cases our weightings were assigned as the inverse of the number of cases which ensures an approximately equal emphasis on AMP and non-AMP cases overall.

With the above considerations in mind we chose a background dataset consisting of 14,830 proteins as follows:

1. Start with all SwissProt proteins clustered to 90% identity
2. Remove any proteins with the keyword “Antimicrobial”
3. Remove very short (<50 AA) and very long (>500 AA) proteins
4. Remove proteins with non-standard amino acids.
5. Randomly sample to obtain 10x the number of AMPs

**Mature Peptide Model Training Data**

For the mature peptide model we selected positive (AMP) cases as follows:

1. Include all AMPs from the APD, DRAMP and dbAMP databases with lengths >10 AA and < 60 AA
2. Include mature peptides from SwissProt (also with length >10 AA and <60 AA)
3. Remove sequences that are identical or that contain non-standard amino acids
4. Cluster sequences to 90% identity using CD-HIT.

A total of 3,232 positive AMP mature peptide sequences were obtained using these criteria and a negative (non-AMP) dataset of equal size was constructed as follows;

1. Start with a complete listing of all proteins in the SwissProt database.
2. Cluster these to 90% identity using CD-HIT
3. Remove sequences with non-standard amino acids
4. Keep only sequences with lengths between 10 and 40

This resulted in a total of 3,321 background mature peptides. Note that the length criterion for background peptides was stricter than for the target set because the target set is more carefully curated and much less likely to include short precursor sequences. We found that a large number of sequences around 50-60 AA in the background data were likely precursors and these were effectively filtered using a length cut-off of 40 AA.

#### Feature Calculation and Selection

ampir uses a suite of features commonly used in AMP prediction, including the physicochemical properties, amphiphilicity, charge, hydrophobicity, molecular weight and pI as well as a further 39 features known as Chou's pseudo amino acid composition (Xiao *et al.*, 2013). All features were centred and scaled for model fitting and prediction.

In order to select features for the default ampir models we examined each feature individually to determine whether its values were related to the model classes (AMP, Non-AMP). We did this through visual inspection of the probability densities for background and target sets based on the training data (see https://github.com/Legana/AMP_pub/blob/master/03_feature_selection.md) for actual plots used. All features showing a clear separation between AMP and Non-AMPs were included. On this basis we constructed default models using all physicochemical properties as well as all Chou's pseudo-amino acid composition terms except for lambda values higher than two. Note that the reserved test data was not used in feature selection.

#### Model Training and Tuning

ampir uses the support vector machine with radial kernel for its default models but accepts many other machine learning predictors supported by the caret package (Kuhn, 2008) as custom models.

For the purposes of model tuning and testing the same procedure was used for both mature and precursor models. Firstly, the full training dataset was split into a training component (80%) and a hold-out testing (20%) set. The caret package was then used to train an SVM with radial kernel with the training data. The parameters sigma and C were tuned by searching a grid of values and choosing those with the highest accuracy as determined by 3 repeats of 10-fold cross validation.

The 20% hold-out test set was used to evaluate ampir as well as other predictors (see below). For the final model distributed with the ampir package this evaluation set was combined with the initial training set to train a final version of the model for distribution.

### Section S4 Model performance evaluation on various benchmark sets

**Table S1: Performance of ampir and other AMP predictors on three different test sets**

| Metric | ampir_mature | ampir_precursor | ampscanner v2 | amPEP | iAMPpred |
|---|---|---|---|---|---|
| ***Xiao et al. 2013 benchmark set*** | | | | | |
| Accuracy | 0.97 | 0.61 | 0.83 | 1.00 | 0.64 |

| | | | | | |
|---|---|---|---|---|---|
| F1 score | 0.97 | 0.44 | 0.85 | 1.00 | 0.73 |
| AUROC | 0.99 | 0.80 | 0.94 | 1.00 | 0.86 |
| ***ampir_mature test set*** | | | | | |
| Accuracy | 0.91 | 0.58 | 0.73 | 0.76 | 0.70 |
| F1 score | 0.91 | 0.39 | 0.56 | 0.58 | 0.53 |
| AUROC | 0.97 | 0.66 | 0.79 | 0.90 | 0.74 |
| ***ampir_precursor test set*** | | | | | |
| Accuracy | 0.50 | 0.94 | 0.70 | 0.46 | 0.47 |
| F1 score | 0.17 | 0.92 | 0.26 | 0.06 | 0.16 |
| AUROC | 0.84 | 1.00 | 0.82 | 0.52 | 0.50 |

AUROC: area under the receiver operating characteristics curve

**Table S2: Performance of ampir and other AMP predictors on ampir's training data**

| **Metric** | **ampir_mature** | **ampir_precursor** | **ampscanner v2** | **amPEP** |
|---|---|---|---|---|
| ***ampir_mature training data*** | | | | |
| Accuracy | 0.92 | 0.58 | 0.76 | 0.76 |
| F1 score | 0.92 | 0.35 | 0.79 | 0.80 |
| AUROC | 0.97 | 0.67 | 0.83 | 0.90 |
| ***ampir_precursor training data*** | | | | |
| Accuracy | 0.50 | 0.97 | 0.71 | 0.46 |
| F1 score | 0.17 | 0.96 | 0.27 | 0.06 |
| AUROC | 0.86 | 1.00 | 0.82 | 0.53 |

In Figure S4 (first row) we show how various predictors perform on genome-scanning data as a tradeoff between sensitivity (proportion of AMPs detected among all AMPs present in the dataset) and precision (proportion of true positives compared with all positive results). This shows that in order to obtain a high precision it is necessary to discard many true AMPs (low sensitivity). Note that the ampir_precursor model clearly outperforms all other models in this context.

In Figure S4 (second row) we show ROC curves for all predictors on the Human and *Arabidopsis thaliana* data but it is important to remember that in this context the low false positive regime is especially important. This is because of the extremely low frequency of true positives in the data (less than 1%). This is explored further in Figure 1 but for now it is important to note that `ampscanner v2` is not shown in Figure 1 (A and B) because its ROC curve does not extend into this important regime despite the fact that it otherwise appears to perform very well. `AmPEP` and `ampir_mature` both perform very poorly reflecting the emphasis of their training data on mature peptides rather than precursor proteins.

In order to properly capture the real-world performance of predictors on genome scans we use a plot that emphasises the absolute numbers of true and false positives. On this measure (shown in Figure 1) it can be seen that genome-wide prediction of AMPs is still an imperfectly solved problem. Although the `ampir` precursor model clearly performs far better than any other predictors, none were able to predict more than 50% of true AMPs while controlling false positives to under 500. Nevertheless, given the difficulties in identifying AMPs and the importance of this task this level of enrichment is of great practical use, reducing the number of false experimental leads per true positive from many thousands down to tens or hundreds.

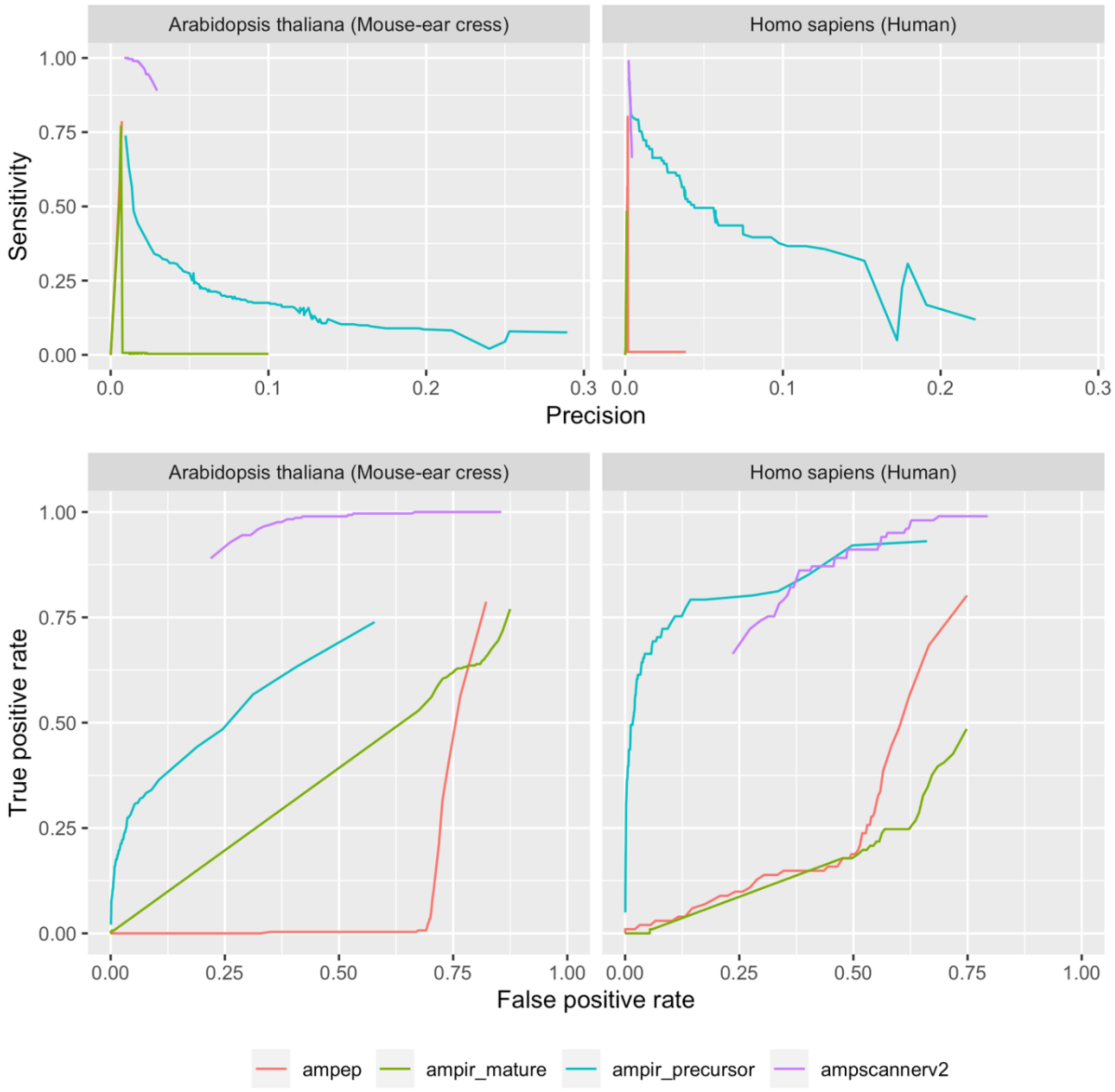


**Figure S4: Performance of various AMP predictors in classifying whole proteome data for Human and Arabidopsis thaliana. Performance is shown as a balance of sensitivity and precision (top row) and as a ROC curve (second row).**

### Section S5 Running time

Benchmarking the computational speed of AMP predictors is difficult to do in a completely objective fashion. This is because many predictors are available only through web servers, in which case the performance depends on unknown factors such as server load and configuration. Here we present some approximate benchmarks for the speed of ampir, ampscanner v2 and ampep based on a complete Human proteome dataset (74,811 proteins) (see Table S2). Performance of iamppred could not be evaluated with this dataset as we found that it was unable to handle large numbers of input sequences.

*ampscanner v2:*
Maximum file upload size is 50Mb so it was necessary to divide the job into two parts. A stopwatch was used to measure runtime. Timing was started after the upload step had finished so as not to include internet connectivity speed in the test. The reported run time is the sum of runtimes for both parts of the dataset.

*ampep*:
ampep was run using MATLAB R2019a on an Intel Xeon processor with 40 CPUs but runtime reflects single core performance since ampep did not appear to have a multi-core capability.

*ampir:*
ampir was run using the same Intel Xeon processor as was used for the `ampep` benchmark. Since ampir is capable of multicore operation we measured its runtime as a function of core-count (see Figure 5.4). In Table S2 the runtime of ampir with a single core is shown. ampir provides comparable performance to ampscanner v2 when run with four cores.

**Table S2: Run time performance on the Human Proteome**

| Program | Number of cores | Runtime (s) |
|---|---|---|
| ampscanner v2 | Unknown | 195 |
| AmPEP | 1 | 1223 |
| ampir | 1 | 469 |
| ampir | 4 | 218 |
| ampir | 8 | 110 |

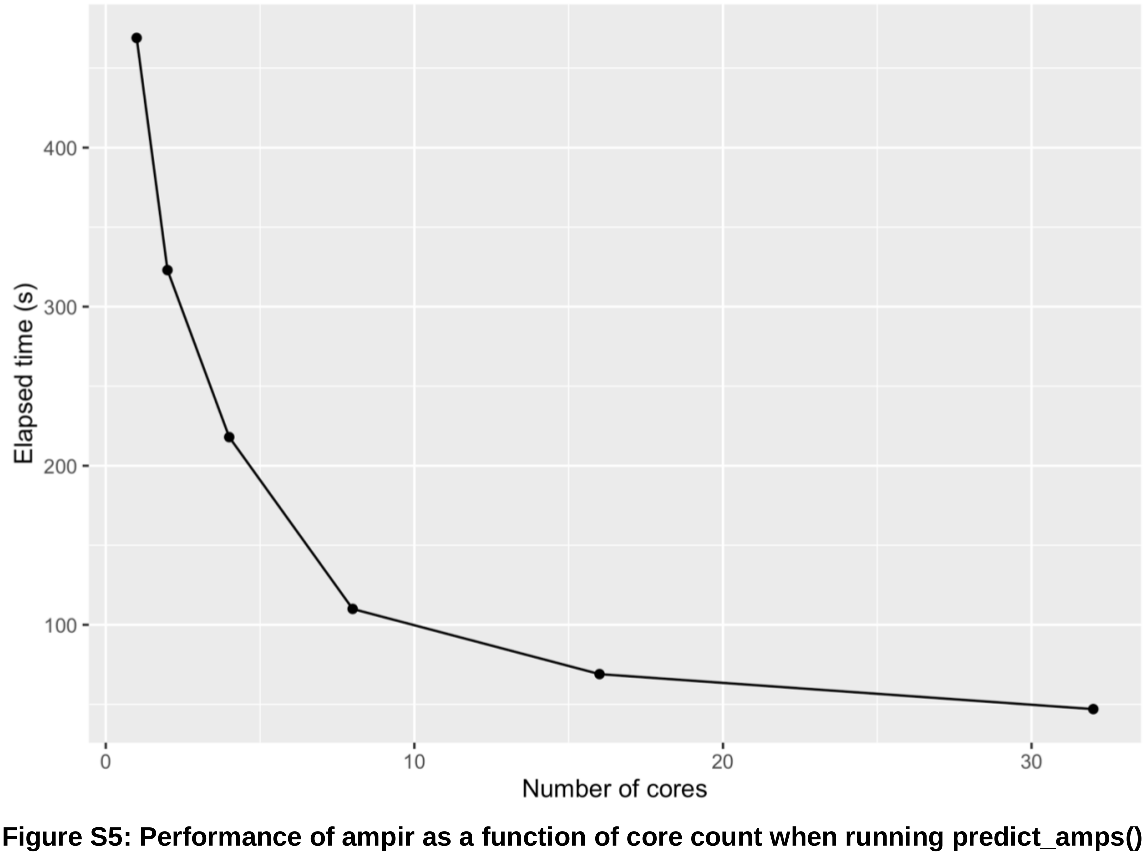


**Figure S5: Performance of ampir as a function of core count when running predict_amps() on a dataset of 77,000 proteins**
